# Collagen-binding by *Drosophila* SPARC is essential for survival and for collagen IV distribution and assembly into basement membranes

**DOI:** 10.1101/714378

**Authors:** Sebastian Duncan, Samuel Delage, Alexa Chioran, Olga Sirbu, Theodore J. Brown, Maurice J. Ringuette

**Author notes:** Contributed equally to the work, co-first authors (Sebastian Duncan, Samuel Delage). Address Correspondence to: Dr. Maurice J. Ringuette, Department of Cell and Systems Biology, University of Toronto, 25 Harbord Street, Toronto, Ontario, Canada M5S 3G5.

## Abstract

The assembly of basement membranes (BMs) into tissue-specific morphoregulatory structures requires non-core BM components. Work in *Drosophila* indicates a principal role of collagen-binding matricellular glycoprotein SPARC (Secreted Protein, Acidic, Rich in Cysteine) in larval fat body BM assembly. We report that SPARC and collagen IV (Col(IV)) first colocalize in the trans-Golgi of hemocytes. Mutating the collagen-binding epitopes of SPARC leads to 2^nd^ instar larval lethality, indicating that SPARC binding to Col(IV) is essential for survival. Analysis of this mutant reveals increased Col(IV) puncta within adipocytes and intense perimeter Col(IV) staining surrounding the fat body as compared to wild-type larvae, reflecting a disruption in chaperone-like activity. In addition, Col(IV) in the wing imaginal disc was absent. Removal of the disulfide bridge in EF-hand2, which is known to enhance Col(IV) binding by SPARC, did not lead to larval lethality; however, a similar but less intense fat body phenotype was observed. Additionally, both SPARC mutants have altered fat body BM pore topography. Wing imaginal disc-derived SPARC did not localize within Col(IV)-rich matrices, indicating a distinct variant. Collectively, these data demonstrate the essential role of Col(IV) chaperone-like activity of SPARC to *Drosophila* development and indicate tissue-specific variants with differential functions.

## Introduction

Basement membranes (BMs) are specialized sheet-like extracellular matrix networks with tissue-specific morphoregulatory and physiological functions. Context-dependent functions include tissue epithelization and compartmentalization, cell adhesion and migration, establishment of growth factor gradients, and creation and maintenance of stem cell niches (Adams, 2018; Arnaoutova et al., 2012; Isabella and Horne-Badovinac, 2015a; Pozzi et al., 2017). Tissue-specific adaptations in the BM affecting thickness, stability, tensile strength, and porosity enable specialized organ function. As such, BMs play an indispensable role in development, growth, regenerative capacity and tissue homeostasis. Consequently, mutations or dysregulation in the expression, assembly, and remodeling of BM constituents is a major contributor to the underlying cause of numerous pathologies (Wiradjaja et al., 2010).

BMs are comprised of four universal core components: laminins, collagen IV (Col(IV)), perlecan and nidogens. The diversity in the structure and functions of BMs is only partly attributable to variability within the laminin and Col(IV) family members. Emerging evidence indicates that tissue-specific non-core components make important contributions to BM assembly and remodeling (Randles et al., 2017).

The initial step in the assembly of all BMs involves laminin anchoring and polymerization at cell surface receptors, most notably dystroglycans and integrins. Col(IV), which imparts tensile strength to the BM, then assembles as a second network, anchoring to the laminin network predominantly through the proteoglycans and nidogens. Col(IV) protomers are composed of three α-chains that assemble into three structural domains: a long N-terminal 7S domain, a central discontinuous collagenous domain, and a C-terminal globular domain (NC1). The first step in Col(IV) polymerization, which may occur prior to secretion, is the formation of C-terminal dimers (Chioran et al., 2017; Yurchenco et al., 2004) These dimers are deposited and begin to polymerize onto the laminin network, where they begin to form 7S anti-parallel tetramers that are stabilized by disulfide bonds and lysine and hydroxylysine cross-links. Electron microscopy rotary shadowing analyses indicate that the stability of the Col(IV) networks are enhanced by lateral associations within the flexible discontinuous collagenous domains (Eble et al., 1996).

Since Col(IV) has an innate ability to self-polymerize (Chioran et al., 2017), developmental strategies must exist to prevent both intracellular polymerization and premature extracellular Col(IV) polymerization. This is particularly important for invertebrates where Col(IV) and other core components are synthesized and must diffuse to and assemble into BMs within distal tissues. In vertebrates, Hsp47 has been identified as a collagen-specific chaperone that ensures proper Col(IV) protomer assembly in the endoplasmic reticulum (ER) and transit to the cis-Golgi while preventing aberrant aggregation; however, Hsp47 is not expressed in invertebrates. It therefore remains to be elucidated how Col(IV) polymerization is prevented in invertebrates during its transit from the ER to the Golgi network. Moreover, for these organisms, it is critical that Col(IV) polymerization be delayed until reaching its final destination, often distal to its source of synthesis.

Studies in *D. melanogaster* and *C. elegans* report that the proper assembly of Col(IV) into BMs is dependent on the presence of SPARC (Secreted Protein Acidic Rich in Cysteine). Moreover, the diffusion of Col(IV) to secondary sites is inhibited by the absence of SPARC (Isabella and Horne-Badovinac, 2015b; Morrissey et al., 2016; Pastor-Pareja and Xu, 2011; Shahab et al., 2015). It is hypothesized that SPARC functions to delay nucleation of Col(IV) polymers upon their secretion to enable proper BM assembly at both the site of production and at distal sites that do not express BM components. Based on this model, SPARC is predicted to associate with Col(IV) prior to secretion to prevent precocious Col(IV) polymerization. Colocalization of SPARC and Col(IV) in numerous punctate structures is observed in *Drosophila* hemocytes (Martinek et al., 2008). However, the point at which SPARC associates with Col(IV) in the secretory process remains to be determined.

SPARC is composed of three structural domains: an acidic Ca^2+^-binding N-terminal domain, a central follistatin-like domain, and a C-terminal domain containing two highly conserved collagen-binding epitopes and two high-affinity Ca^2+^-binding EF-hands. These collagen-binding epitopes and EF-hands of SPARC are highly conserved in animals ranging from cnidarians to mammals. The affinity of SPARC for collagens is enhanced by the cooperative interaction of the collagen-binding epitopes with the EF-hands (Pottgiesser et al., 1994). The principle source of core BM components in *Drosophila* larvae is the fat body (Chioran et al., 2017; Pastor-Pareja and Xu, 2011). The loss of SPARC leads to 2^nd^ instar larval lethality, characterized by an aberrant BM accumulation in the fat body and an absence of Col(IV) diffusion to distal tissues (Shahab et al., 2015).

In this study, we define the precise stage at which SPARC and Col(IV) colocalize during the secretory pathway. Moreover, we establish the importance of the collagen-binding epitopes and the structural integrity of the disulfide bridge-dependent EF-hand2 to the Col(IV) chaperone-like activity of SPARC. We also provide evidence that suggests larval fat body-derived SPARC is functionally distinct of that produced by the wing imaginal disc and reveal a potential mechanism to explain the 2^nd^ instar lethality observed with the loss of SPARC.

## Results

### Intracellular colocalization of SPARC with Col(IV) is first evident in the trans-Golgi network of cultured hemocytes

*Drosophila* embryonic hemocyte-like cell lines (S2 and Kc167), which express endogenous SPARC and Col(IV) (Drosophila Genomic Resource Centre, https://dgrc.bio.indiana.edu/cells/Catalog, unpublished data), were used to examine when these two proteins first colocalized within the secretory pathway. Immunocytochemistry performed on S2 and Kc167cells indicate extensive intracellular colocalization of SPARC and Col(IV) (Fig. 1A). A panel of secretory pathway markers (Riedel et al., 2016) was used to track the intracellular distribution of SPARC and Col(IV) in the secretory pathway. An antibody against calnexin 99A, an ER-specific folding chaperone protein, was used to visualize the ER (Riedel et al., 2016). Immunostaining of S2 cells with anti-calnexin 99A revealed ring-shaped perinuclear structures and discrete puncta, consistent with ER morphology and the extensive nature of the ER lumen of *Drosophila*-cultured hemocytes (Zacharogianni and Rabouille, 2013) (Fig. 1B). Small puncta positive for SPARC colocalized with calnexin 99A within the perinuclear region. Less extensive punctate staining was observed for SPARC than calnexin 99A. However, punctate SPARC staining was also observed throughout the cytoplasm that did not colocalize with calnexin 99A. In contrast, Col(IV) foci did not appear to localize within ER-labelled structures. Immunostaining for SPARC, Col(IV), and calnexin 99A confirmed that SPARC and Col(IV) did not colocalize within the ER. The lack of Col(IV) staining in the ER was surprising since all secretory proteins, including Col(IV), are translated into the ER where Col(IV) assembly into protomers is tightly regulated by chaperones. The absence in staining may be attributable to epitope masking as a result of the binding of chaperones to the globular NC1 domain of Col(IV) against which the Col(IV) serum was raised (Shahab et al., 2015). Thus, the possibility that SPARC and Col(IV) associate within the ER cannot be completely discounted. This limitation prompted us to investigate whether SPARC and Col(IV) could be detected leaving the ER together.

**Fig 1.**
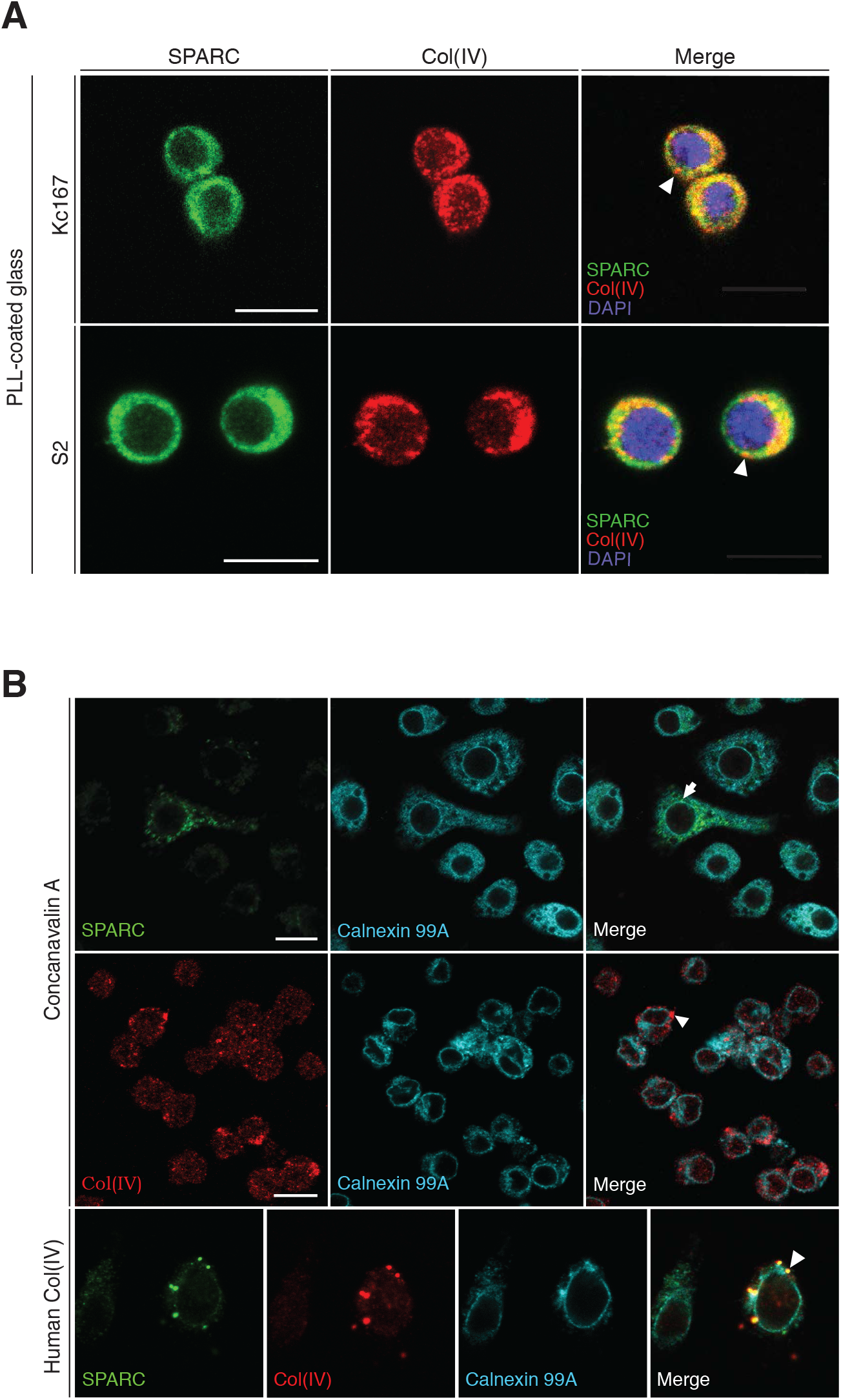
Intracellular distribution of SPARC and Col(IV) within S2 and Kc167 embryonic hemocyte-like cells indicates colocalization in the trans-Golgi. **(A)** Kc167 and S2 cells seeded on PLL-coated glass and stained with anti-SPARC and anti-Col(IV) antibodies. Merged images indicate extensive overlap in the punctate distribution of SPARC and Col(IV). Arrowheads denote that some Col(IV) foci do not colocalize with SPARC. Cells were counterstained with DAPI to mark chromatin. **(B)** S2 cells seeded on Con-A and stained with anti-SPARC, anti-Col(IV) and anti-calnexin 99A antibodies, the latter to mark the ER. The arrow denotes SPARC and calnexin 99A immunostaining overlaps in the region surrounding the nucleus. In contrast, no immunostaining is evident for Col(IV) within the ER (arrowhead). S2 cells seeded on human Col(IV) showed a similar pattern with SPARC colocalizing with calnexin 99A but not Col(IV). All images were taken at 63X objective. Scale bars represent 10 μm.

Anterograde movement along the secretory pathway is facilitated by the formation of COPII vesicles, which relay cargo from the ER to the cis-Golgi and from the trans-Golgi to the plasma membrane. The COPII-associated protein Tango1 delays the normal budding of COPII vesicles from ER exit sites to accommodate larger proteins, such as Col(IV). An antibody against Tango1 (Lerner et al., 2013) was used as a marker for ER exit sites and vesicles travelling from the ER to the cis-Golgi. Due to the common host species of the anti-Tango1 and anti-Col(IV) antibodies, simultaneous staining of Col(IV) and Tango1 was not performed. However, due to their size, all Col(IV)-containing vesicles would be expected to be Tango1-positive (Liu et al., 2017). Co-immunocytochemical staining for SPARC and Tango1 indicated that SPARC was rarely present in Tango1-positive vesicles (Fig. 2A). Pearson Correlation coefficients indicate that SPARC and Tango1 colocalized (PC=0.30 ±0.03) significantly less than SPARC and Col(IV) (PC=0.77 ±0.02) (Fig. 2B). Thus, it is unlikely that SPARC and Col(IV) exit the ER within the same vesicle. We then explored the possibility that SPARC and Col(IV) are co-packaged within the Golgi Apparatus.

**Fig 2.**
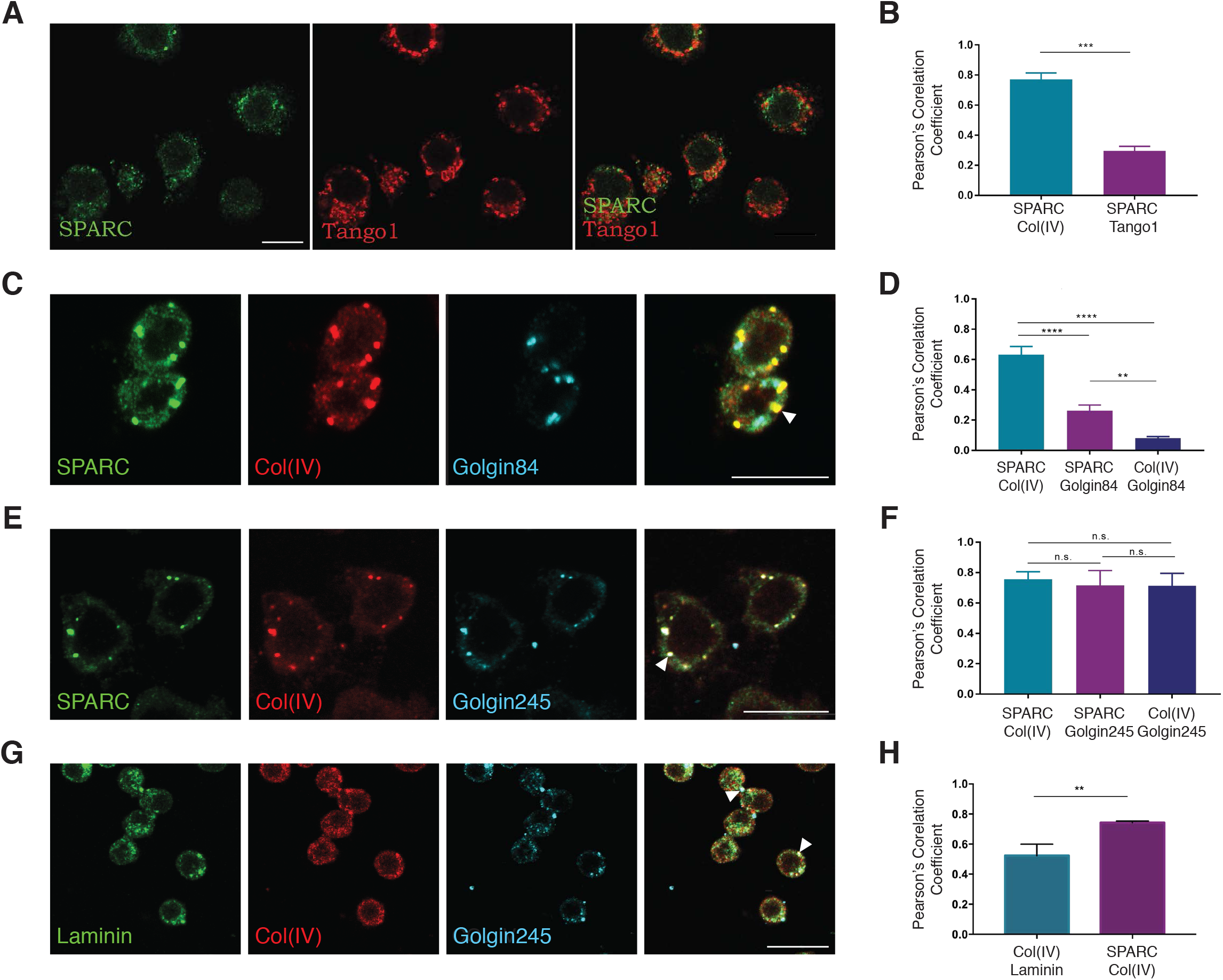
SPARC and Col(IV), but not laminin, colocalize in the trans-Golgi network of the secretory pathway. **(A)** S2 cells seeded on Con-A were stained with anti-SPARC and anti-Tango1 antibodies, the latter to mark ER exit sites and modified ER-to-Golgi transport vesicles. SPARC shows little localization within Tango1-positive vesicles. **(B)** Quantification of SPARC and Tango1 colocalization. The graph shows the mean Pearson Correlation coefficient for Tango1 and SPARC (0.30 ±0.03) calculated for three independent trials (trial 1 n=32, trial 2 n=26, trial 3 n=29). The mean Pearson correlation coefficient for SPARC and Col(IV) (0.77 ±0.02) is calculated from a separate experiment (n=25). Bars represent mean ± SD. An unpaired t-test was used to determine statistical significance. P<0.001 (***). **(C)** S2 cells seeded on Con-A were stained with anti-SPARC, anti-Col(IV), and anti-Golgin84 antibodies, the latter of which marks the medial-Golgi network. SPARC and Col(IV) positive foci, denoted by the arrowhead, are infrequently found within the medial-Golgi. **(D)** Quantification of SPARC and Col(IV) colocalization within the medial-Golgi. SPARC and Col(IV) colocalization is compared to the colocalization of either SPARC or Col(IV) with Golgin84. The graph represents the mean Pearson Correlation coefficient of SPARC and Col(IV) (0.63 ±0.03), SPARC and Golgin84 (0.26 ±0.02), and Col(IV) and Golgin84 (0.08 ±0.01) calculated from three trials (trial 1 n=30, trial 2 n=32, trial 3 n=27). Bars represent mean ± SD. Unpaired t-tests were used to determine statistical significance. P<0.01 (**), P<0.0001 (****). **(E)** S2 cells seeded on Con-A were stained with anti-SPARC, anti-Col(IV), and anti-Golgin245 antibodies, the latter of which marks the trans-Golgi network. SPARC and Col(IV) colocalization is observed in the trans-Golgi network as large white foci indicated by the arrowhead. **(F)** Quantification of SPARC and Col(IV) colocalization within the trans-Golgi network. The graph represents the mean Pearson Correlation coefficient of SPARC and Col(IV) (0.72 ±0.03), SPARC and Golgin245 (0.63 ±0.06), and Col(IV) and Golgin245 (0.65 ±0.05) calculated form three trials (trial 1 n=39, trial 2 n=22, trial 3 n=25). Bars represent mean ± SD. ANOVA followed by multiple t-test comparisons was used to determine significance. P<0.01 (**), P<0.0001 (****), P>0.05 (n.s.). **(G)** S2 cells seeded on Con-A were stained with anti-laminin, anti-Col(IV), and anti-Golgin245 antibodies. Arrowheads indicate the few events of Col(IV) and laminin colocalization. Col(IV) colocalizes with laminin significantly less than with SPARC. **(H)** The graph shows the mean PC coefficient calculated for three independent trials comparing laminin and Col(IV) overlap (0.52 ±0.04, n= 75) and SPARC and Col(IV) overlap (0.74 ±0.01, n=73) in S2 cells. Bars represent mean ± SD. An unpaired t-test was performed to determine significance. P<0.01 (**). All images were taken at 63X objective. Scale bars represent 10 μm.

The medial-Golgi and trans-Golgi were visualized by the transmembrane structural proteins Golgin84 and Golgin245, respectively (Riedel et al., 2016). Pearson Correlation coefficients consistently showed high colocalization of SPARC and Col(IV) overall (PC=0.63 ±0.03). The colocalization of SPARC and Col(IV) was detected in the medial-Golgi network, albeit at low levels (Fig. 2C). The colocalization of either SPARC or Col(IV) with Golgin84 (PC=0.26 ±0.02 and 0.08 ±0.01, respectively) was significantly lower (Fig. 2D), suggesting that the majority of foci positive for both Col(IV) and SPARC did not colocalize within the medial-Golgi.

Strong colocalization of SPARC and Col(IV) became evident in the trans-Golgi network, where foci positive for Golgin245 stained for both SPARC and Col(IV) (Fig. 2E). Pearson Correlation coefficients demonstrate the overlap of either SPARC or Col(IV) with Golgin245 (PC=0.63 ±0.06 and 0.65 ±0.05, respectively) with a comparable overlap between SPARC and Col(IV) (PC=0.72 ±0.03) (Fig. 2F). The strong colocalization of SPARC and Col(IV) in the trans-Golgi network indicates that these proteins are co-packaged just prior to their transport to the plasma membrane for secretion. However, it remains possible that this colocalization may simply reflect a shared destination of cargo rather than a functional interaction. If the latter, then SPARC should colocalize with equal frequency to other core BM components, such as laminin. Our data indicate that neither SPARC nor Col(IV) colocalized with laminin in Golgin245-positive vesicles (Fig. 2G). The mean Pearson Correlation coefficient (0.52 ±0.04) calculated from the overlap of laminin and Col(IV) in cultured hemocyte cells is significantly lower than the mean Pearson Correlation coefficient calculated for SPARC and Col(IV) (0.74 ±0.01) (Fig. 2H).

### Disruption of the SPARC locus using CRISPR-Cas9 results in 2^nd^ instar larval lethality

We previously generated a SPARC deficiency line (*Df(3R)nm136, H2Av*∷GFP) by P-element excision which resulted in 1^st^ instar larval lethality in a homozygous state. Complementation analyses were consistent with a second mutation site unrelated to SPARC, such that in the heterozygous state, lethality occurred at the 2^nd^ instar (Shahab et al., 2015). To generate a mutant fly line specifically targeting SPARC, for further use, the CRISPR-Cas9 approach was used. Two SPARC protein-*null* fly lines, *SPARC*^*1510B*^ and *SPARC*^*1510D*^ were generated. Genomic analyses revealed that these SPARC protein-*null* lines contained identical 2795 bp deletions overlapping the 5’UTR of the SPARC locus and the 3’UTR of the adjacent RB97D gene (Fig. S1). Viability assays indicated that the greatest percentage of larval death occurred at 2^nd^ instar larvae for both *SPARC*^*1510B*^ (67.7% ±2.8) and *SPARC*^*1510D*^ (74.8% ±0.8) homozygotes, consistent with the lethality stage observed in *Df(3R)nm136, H2Av*∷GFP homozygotes. Similarly, heterozygote larvae of *SPARC*^*1510B*^/*Df(3R)nm136, H2Av*∷GFP and *SPARC*^*1510D*^/*Df(3R)nm136, H2Av*∷GFP also exhibited 2^nd^ instar larval lethality, further validating that disruption of SPARC expression is responsible for this lethality.

The unexpected perturbation to the 3’UTR of *Rb97D*, which encodes for a ribonucleic protein, raises the possibility that this contributed to the larval lethality observed in the CRISPR mutant flies. To test for this, SPARC expression was restored in the SPARC mutants by crossing the mutants to flies containing a UAS-SPARC insertion (UAS-SPARC-16) under the control of the endogenous SPARC promoter (MIMIC *SPARC*-Gal4) to mimic the expression profile of SPARC in wild-type flies (Shahab et al., 2015). Transgenic expression of UAS-SPARC-16 with *SPARC*-Gal4 rescued *SPARC*^*1510B*^ and *SPARC*^*1510D*^ mutants to adulthood. Driving the expression of UAS-SPARC-16 with *Cg*-Gal4, which induces expression of SPARC only in hemocytes and the fat body, also rescued both *SPARC*^*1510B*^ and *SPARC*^*1510D*^ homozygous mutants to adulthood, indicating that the deficit of SPARC leads to larval lethality. The *SPARC*^*1510D*^ mutant was used for all subsequent experiments.

We next verified the loss of SPARC protein in the *SPARC*^*1510D*^ mutant and whether it impacted the spatiotemporal distribution of Col(IV) during embryogenesis. As expected, SPARC was expressed by circulating hemocytes and concentrated in the BM of the developing gut and VNC of wild-type embryos, but was absent in the *SPARC*^*1510D*^ mutant. Col(IV) protein distribution mirrored that of SPARC in wild-type embryos. As previously reported, loss of SPARC had no observable impact on the distribution of Col(IV) at this stage of development (Martinek et al., 2008; Shahab et al., 2015) (Fig. S2).

We then examined the distribution of Col(IV) at the 2^nd^ instar stage when larval lethality was observed in *SPARC*^*1510D*^. Mutant larvae showed intense pericellular and intracellular accumulation of Col(IV) in fat body adipocytes compared to a less intense BM localization in wild-type larvae (Fig. 3C).

**Fig 3.**
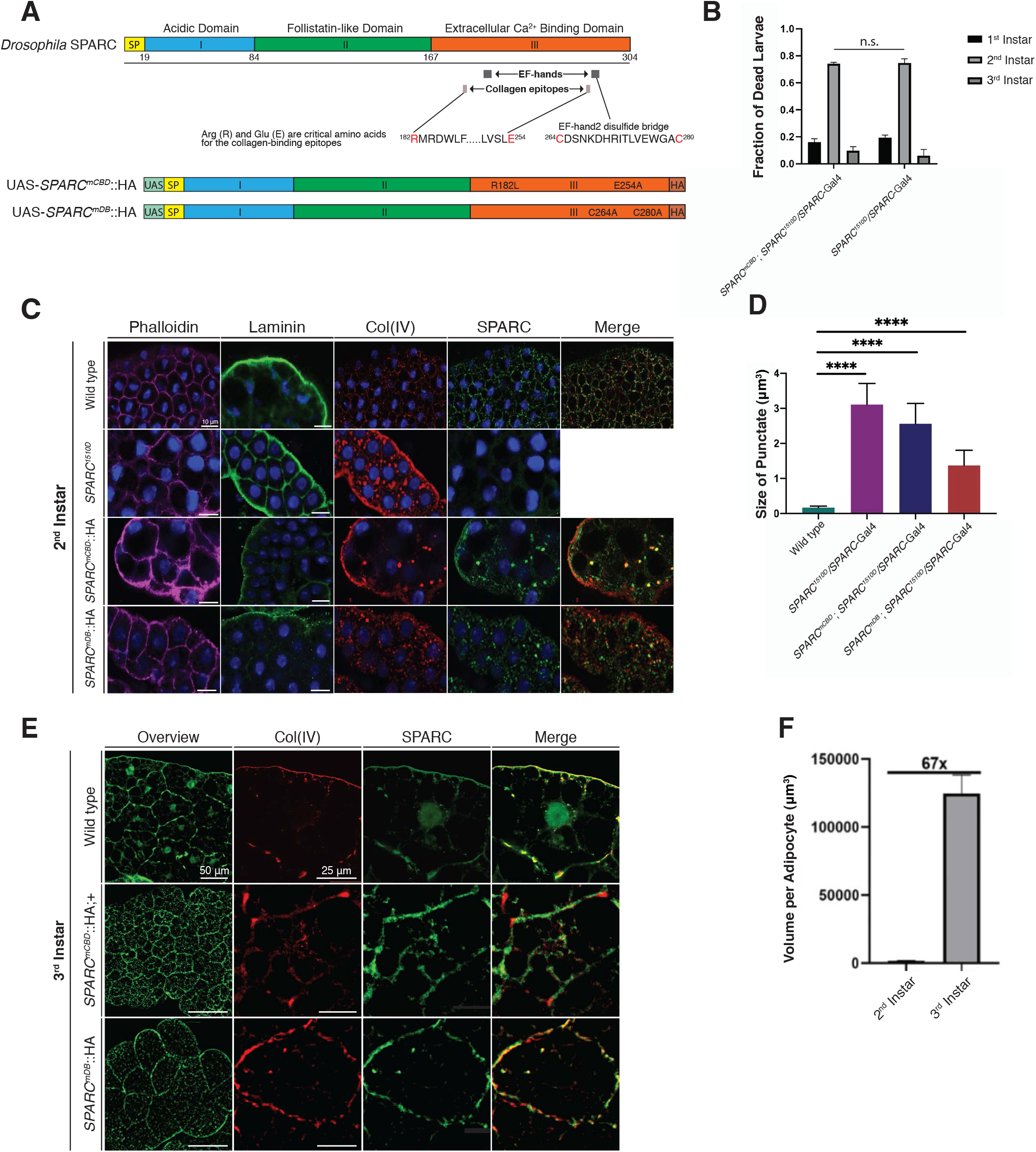
Mutation of SPARC alters colocalization with Col(IV) at 2^nd^ and 3^rd^ instar. **(A)** Schematic representation of transgenic SPARC constructs generated using Gateway Cloning technology. The schematic indicates the UAS promoter (teal), a signal peptide (SP; yellow), SPARC Domain I (blue), SPARC Domain II (green), SPARC Domain III (fuchsia), and C-terminal HA tag (brown). Amino acids critical for collagen binding and cysteines involved in the formation of the disulfide bridge in EF-hand2 are highlighted in red. The amino acids substitutions for the conserved collagen-binding epitopes (UAS-*SPARC*^*mCBD*^∷HA) and cysteine residues (UAS-*SPARC*^*mCBD*^∷HA) are indicated within Domain III of both constructs. **(B)** Larval lethality for *SPARC*-Gal4/*SPARC*^*1510D*^ and *SPARC*^*1510D*^/*SPARC*-Gal4>*SPARC*^*mCBD*^∷HA shows lethality occurring at 2^nd^ instar with no statistical significance between the two genotypes. Data were analyzed by one-way ANOVA followed by Dunnett’s multiple comparisons test, P>0.05 (n.s.) **(C)** Immunohistochemical analysis of 2^nd^ instar fat body for wild-type SPARC, *SPARC*^*1510D*^/*SPARC*-Gal4, *SPARC*^*mDB*^∷HA ; *SPARC*^*1510D*^/*SPARC*-Gal4, and *SPARC*^*mCBD*^∷HA ; *SPARC*^*1510D*^/*SPARC*-Gal4. Larvae were immunostained with anti-laminin, anti-Col(IV), and anti-SPARC and were stained for actin (phalloidin). Laminin was performed in a parallel experiment due to SPARC and laminin antibody cross-reactivity. As previously shown *SPARC*^*1510D*^ reflects an absence of SPARC protein (Fig. S2). Scale bars indicate 10 μm. **(D)** Size of punctae observed in 2^nd^ instar adipocytes for *SPARC*^*1510D*^/*SPARC*-Gal4, *SPARC*^*mCBD*^∷HA ; *SPARC*^*1510D*^/*SPARC*-Gal4, *SPARC*^*mDB*^∷HA ; *SPARC*-Gal4/*SPARC*^1510D^ relative to wild-type. Data were analyzed by one-way ANOVA followed by Dunnett’s multiple comparisons test, P<0.0001 (****). **(E)** Immunohistochemical analysis of 3^rd^ instar fat body for wild-type SPARC, *SPARC*^*mDB*^∷HA ; *SPARC*^*1510D*^/*SPARC*-Gal4, and *SPARC*^*mCBD*^∷HA ; *SPARC*^*1510D*^/*SPARC*-Gal4. Larvae were immunostained with anti-Col(IV) and anti-SPARC. The overview highlights changes in the size and morphology of adipocytes associated with *SPARC*^*mCBD*^∷HA ; *SPARC*^*1510D*^/*SPARC*-Gal4, *SPARC*^*mDB*^∷HA ; *SPARC*-Gal4/*SPARC*^1510D^ relative to wild-type. *SPARC*^*1510D*^/*SPARC*-Gal4 is not shown due to 2^nd^ instar lethality. Size of scale bars are as indicated in the images. **(F)** Comparison of adipocyte volume at early 2^nd^ instar and late 3^rd^ instar of wild-type larvae.

### Disrupting the collagen-binding capacity of SPARC dysregulates Col(IV) transport and assembly into BMs

The collagen-binding epitopes of SPARC mediate the association of SPARC to Col(IV), which is enhanced by the presence of the disulfide-bridged EF-hand2. We examined the impact of disrupting the capacity of SPARC to bind to Col(IV) by mutating the two collagen-binding epitopes (Sasaki et al., 1998). In addition, we tested the contribution of EF-hand2 alone by removing the disulfide bridge, which is essential for the *in vivo* function of this EF-hand (Pottgiesser et al., 1994).

To facilitate the visualization of the various SPARC constructs, we introduced a C-terminal HA-tag onto UAS-SPARC. This yielded a functional, non-degraded fusion product that was able to rescue the larval lethality in homozygous SPARC protein-null flies. Thus, UAS-*SPARC*∷HA constructs were used in conjunction with our *SPARC*^*1510D*^ protein-null mutant to conduct structure-function studies.

Amino acids critical for the binding of collagen, Arg182 and Glu254 within the collagen-binding epitopes 1 and 2 were substituted with leucine, ^Arg^182^Leu^, and alanine, ^Glu^254^Ala^ (UAS-*SPARC*^*mCBD*^∷HA), respectively (Fig. 3A). Previous studies have shown that these substitutions eliminate Col(IV) binding by SPARC (Sasaki et al., 1998). To disrupt the disulfide bridge of EF-hand2, two cysteine residues were substituted with alanine, ^Cys^264^Ala^ and ^Cys^280^Ala^ (UAS-*SPARC*^*mDB*^∷HA) (Fig. 3A). We examined the impact of these mutations on larval lethality and on Col(IV) fat body distribution and incorporation into BMs.

Transgenic expression of UAS-*SPARC*^*mCBD*^∷HA; *SPARC*^*1510D*^/SPARC-Gal4 did not rescue the larval lethality associated with the lack of SPARC (Fig. 3B), indicating the importance of Col(IV) binding by SPARC for survival. In contrast, transgenic expression of UAS-*SPARC*^*mDB*^∷HA; *SPARC*^*1510D*^/*SPARC*-Gal4 fully rescued larval lethality, suggesting that the enhancement of Col(IV) binding is not essential for survival.

Immunofluorescence and scanning electron microscopy (SEM) were used to determine the impact of the mutations on Col(IV) dynamics and fat body morphology. SPARC and Col(IV) immunofluorescence staining showed substantial punctate colocalization within the cytoplasm and at cell borders of 2^nd^ instar wild-type larval adipocytes. Laminin staining was largely restricted to the edges of the fat body with little internal staining (Fig. 3C). In contrast to wild-type adipocytes (0.17 μm^3^ ±0.02), SPARC protein-null larvae (*SPARC^1510D^/SPARC*-Gal4) Col(IV) puncta were significantly larger and more variable in size (3.11 μm^3^ ±0.30) with greater individual adipocyte cell border staining (Fig. 3C-D). Punctate laminin staining was observed throughout the cytoplasm of the adipocytes rather than being concentrated at the perimeter of the fat body. SEM photomicrographs of wild-type 2^nd^ instar fat body showed that the surface of the polygonal adipocytes were marked with fibrous-like structures with numerous unevenly distributed small pores (Fig. 4A-A’). In the absence of SPARC, fat body adipocytes had a rounded appearance with a fibrous-like ECM coating. The size of the pores was increased 3-fold (0.92 μm ±0.16 *vs.* 0.32 μm ±0.05) while their number decreased by half (212 pores per mm^2^ ±44 *vs.* 417 pores per mm^2^ ±24) (Fig. 4J, 4K, Fig. S4). Knockdown of SPARC resulted in the surface of the fat body being punctuated with numerous pores that have elevated circumferential borders of varying size, which phenocopy the appearance of 3^rd^ instar fat body adipocytes associated with the knockdown of SPARC (Shahab et al., 2015). In 2^nd^ instar fat body, knockdown of SPARC decreased the number of pores per unit area (236 pores per mm^2^ ±41 *vs.* 417 pores per mm^2^ ±24) and increased the size of pores (1.36μm ±0.21 *vs.* 0.32 μm ±0.05) (Fig. 4D-D’, 4H, 4J).

**Fig 4.**
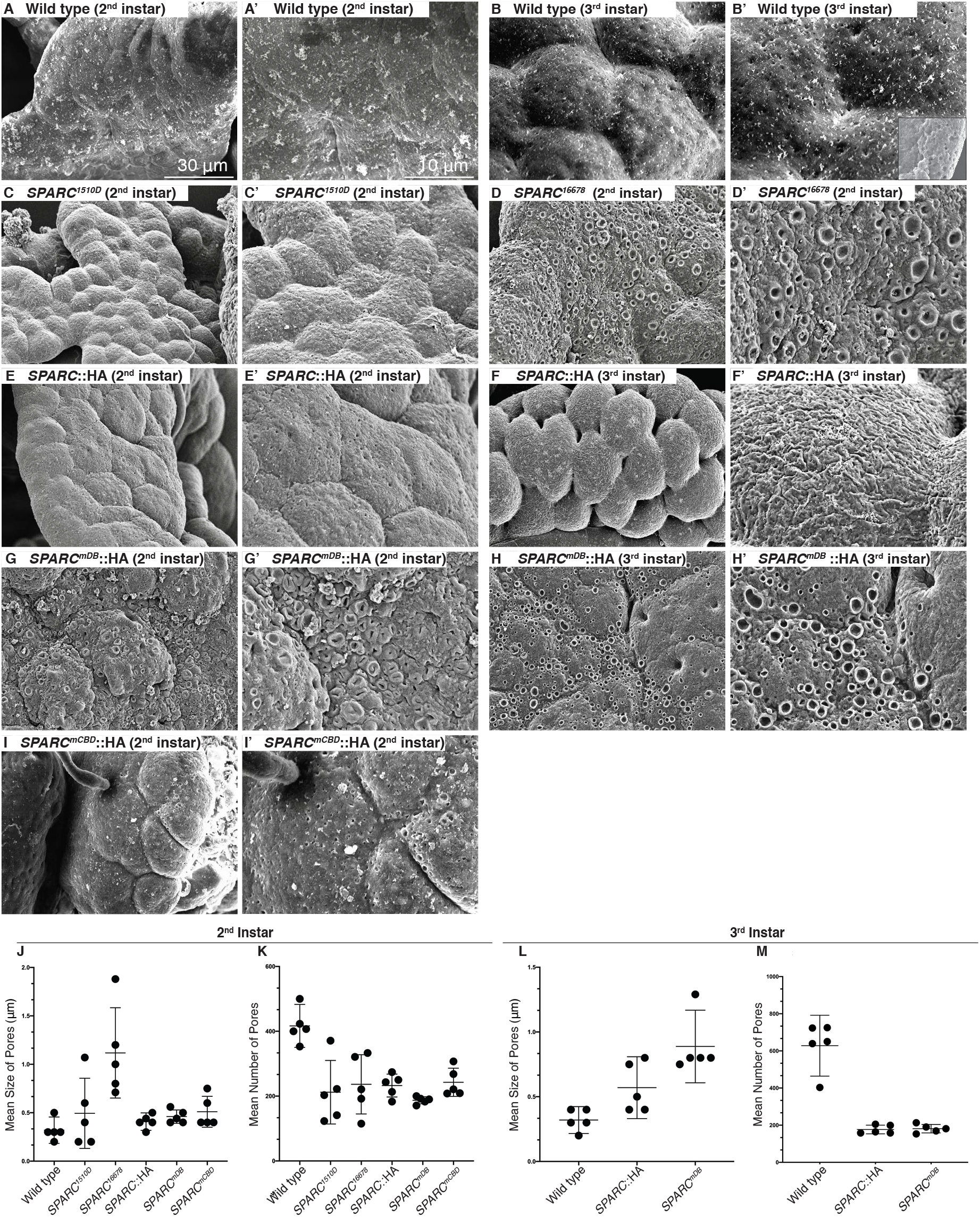
Mutation of SPARC alters the surface topography of 2^nd^ and 3^rd^ instar fat body adipocytes as determined by ultrastructural analysis. SEM of 2^nd^ instar wild-type **(A)**, *SPARC*^*1510D*^ **(C)**, SPARC RNAi (*SPARC*^*16678*^) **(D)**, *SPARC*∷HA **(E)**, *SPARC*^*mDB*^ (**G**), *SPARC*^*mCBD*^ **(I)** fat body at 1700X. **(A’, C’, D’, E’, G’, I’)** Corresponding images at 3700X. SEM of 3^rd^ instar wild-type **(B)**, *SPARC*∷HA **(F)**, *SPARC*^*mDB*^ **(H)** fat body at 1700X. **(B’, F’, H’)** Corresponding images at 3700X. Arrowheads indicate pores. Comparison of the pore size on the surface of 2^nd^ **(J)** and 3^rd^ **(L)** instar fat body adipocytes. Comparison of the number of pores on the surface of 2^nd^ **(K)** and 3^rd^ **(L)** instar fat body adipocytes. Data were analyzed by one-way ANOVA followed by Dunnett’s multiple comparisons test, P<0.0001 (****). Groups with different letters are statistically different from one another, P<0.05.

In contrast to the accumulation of Col(IV) in SPARC protein-null larvae, overexpression of SPARC in a wild-type background resulted in a near absence of Col(IV) within the fat body. Consistent with a previous study (Dai et al., 2017), this loss of Col(IV) in the BM surrounding the fat body was observed in 3^rd^ instar larvae. In addition, individual adipocytes within the fat body were rounded and partially dissociated, as shown by SEM (Fig. 4F-F’). Fat body adipocyte cell-cell adhesion is promoted by Col(IV) Intercellular Concentrations (CIVICs) (Dai et al., 2017). However, our immunohistochemical analyses performed on early 2^nd^ instar wild-type larvae revealed that these CIVICs have not yet formed (Fig. 3C). Despite this, these cells have assumed a polygonal morphology, which is observed in both 2^nd^ instar wild-type and SPARC overexpressed larvae (Fig. 4A-A’, 4E-E’). Immunohistochemical analyses of 3^rd^ instar larvae indicate that CIVICs were present in wild-type larvae but not in SPARC overexpressing larvae, suggesting these CIVICs are required for cell-cell adhesion at the 3^rd^ instar stage but not at 2^nd^ instar. Further SEM analyses of 3^rd^ instar larvae overexpressing SPARC revealed a fibrous-like matrix covering the fat body and extending between adjacent adipocytes, which was not observed in the wild-type fat body (Fig. 4F’). This fibrous-like matrix is likely composed of laminin as perlecan has also been reported to decrease in the fat body of SPARC overexpressing 3^rd^ instar larvae (Isabella and Horne-Badovinac, 2015b). The overexpression of SPARC resulted in increased size and decreased number of pores within the BM at both the 2^nd^ (0.41 μm ±0.03, 233 pores per mm^2^ ±16 *vs.* 0.32 μm ±0.05, 417 pores per mm^2^ ±24) and 3^rd^ instar (0.57 μm ±0.09, 176 pores per mm^2^ ±10 *vs.* 0.32 ±0.04, 628 ±59) stages relative to wild-type larvae (Fig. 4E-E’, 4F-F’, 4J-M, Fig. S4).

Mutating the two collagen-binding epitopes of SPARC resulted in a loss of SPARC colocalization with Col(IV) in the BM. Similar to SPARC protein-null flies, mutation of these epitopes resulted in larval lethality at the 2^nd^ instar stage (Fig. 3B). Immunohistochemical analyses indicated that despite the loss of binding, SPARC and Col(IV) colocalized within punctate structures of variable size (2.56 μm^3^ ±0.29 *vs.* 0.17 μm^3^ ±0.02), indicating that this binding was not required for the intracellular colocalization (Fig. 3D). Importantly, colocalization was not observed extracellularly and Col(IV) was retained intracellularly, demonstrating that SPARC is required for Col(IV) secretion at this stage of development. This entrapment appears to be specific to Col(IV) as laminin did not localize to these large punctate structures. Col(IV) and laminin both appear to be present surrounding the fat body (Fig. 3C). Ultrastructural analyses showed 2^nd^ instar *SPARC*^*mCBD*^∷HA have topographical features similar to 2^nd^ instar *SPARC*^*16678*^, but less pronounced. Some loss of cell-cell adhesion was observed with fewer randomly distributed pores relative to wild-type (243 pores per mm^2^ ±19 *vs.* 417 pores per mm^2^ ±24) The average pore size increased compared to wild-type (1.38 μm ±0.07 *vs.* 0.32 μm ±0.05) (Fig. 4I-I’, 4J, 4M, Fig. S4).

Removal of the disulfide bridge in EF-hand2 (*SPARC*^*mDB*^∷HA) had no impact on *Drosophila* survival. Moreover, *SPARC*^*mDB*^∷HA was able to rescue the larval lethality in a SPARC protein-null background. It was therefore anticipated that loss of the disulfide-bridged EF-hand2 would have less of an impact on fat body morphology and BM homeostasis as compared to disrupting the collagen-binding epitopes. Immunohistochemical analyses of 2^nd^ instar larvae revealed that SPARC and Col(IV) colocalized and were mainly concentrated within the cytoplasm of adipocytes. Intracellular Col(IV) punctae were larger in size compared to wild-type larvae (0.17 μm^3^ ±0.02 vs. 0.17 μm^3^ ±0.02). Laminin immunostaining in wild-type 2^nd^ instar larvae was considerably more restricted to the edge of the fat body than SPARC and Col(IV) (Fig. 3C). Ultrastructural analyses revealed a striking difference in the surface topology of *SPARC*^*mDB*^∷HA 2^nd^ instar larval fat body adipocytes compared to wild-type. The BM of the adipocytes contained numerous variable-sized pores, which were similar in appearance to the knockdown of SPARC (Fig. 4G-G’, 4H-H’). Wild-type pores were on average 0.32 μm (±0.05) in diameter compared to an average pore size of 2.02 μm (±0.03) for *SPARC*^*mDB*^∷HA. Relative to wild-type (417 pores per mm^2^ ±24), *SPARC*^*mDB*^∷HA fat body BM had fewer pores (187 pores per mm^2^ ±5) (Fig. 4J-K, Fig. S4). The surface topography of the BM for *SPARC*^*mDB*^∷HA 2^nd^ and 3^rd^ instar body adipocytes were similar in appearance. No significant differences were observed for the latter in the size or number of pores (Fig. 4J-M, Fig. S4).

As homozygous *SPARC*^*mCBD*^∷HA is lethal at the 2^nd^ instar, *SPARC*^*mCBD*^∷HA larvae were rescued by crossing *SPARC*^*mCBD*^∷HA with the 3^rd^ chromosome *SPARC*-Gal4 driver line, which carries a functional copy of SPARC, to allow immunohistochemical analyses of the impact at the 3^rd^ instar. SPARC was concentrated at adipocyte cell borders of 3^rd^ instar fat body adipocytes in wild-type, *SPARC*^*mDB*^∷HA, and *SPARC*^*mCBD*^∷HA. Moreover, *SPARC*^*mCBD*^∷HA ; *SPARC*-Gal4 fat body adipocytes appeared smaller in size compared to wild-type (Fig. 3E). Under higher magnifications, SPARC and Col(IV) colocalization appeared less evident in *SPARC*^*mCBD*^∷HA ; *SPARC*-Gal4 compared to wild-type and *SPARC*^*mDB*^∷HA, demonstrating haploinsufficiency for *SPARC*^*mCBD*^∷HA.

### Wing imaginal disc-derived SPARC does not associate with BMs

A previous study found that disrupting fat body expression of SPARC resulted in an absence of Col(IV) within the BM of the wing imaginal disc (Pastor-Pareja and Xu, 2011). Similarly, *SPARC*∷HA driven by *Cg*-Gal4, a fat body driver, associated with the fat body BM and diffused to the wing imaginal disc BM (Fig. 5). These data support the model that SPARC functions as an intracellular and extracellular chaperone for Col(IV) (Chioran et al., 2017). Disruption of the collagen-binding epitopes in fat body-derived SPARC resulted in a marked decrease in Col(IV) and a near absence of SPARC within the wing imaginal disc BM (Fig. 5), providing additional support of a chaperone function for SPARC and highlighting the importance of the collagen-binding epitopes. Disruption of the disulfide bridge of EF-hand2 had little to no effect on the diffusion of SPARC and Col(IV) to the wing imaginal disc, indicating that the structural integrity of EF-hand2 is not critical for the chaperone-like activity.

**Fig 5.**
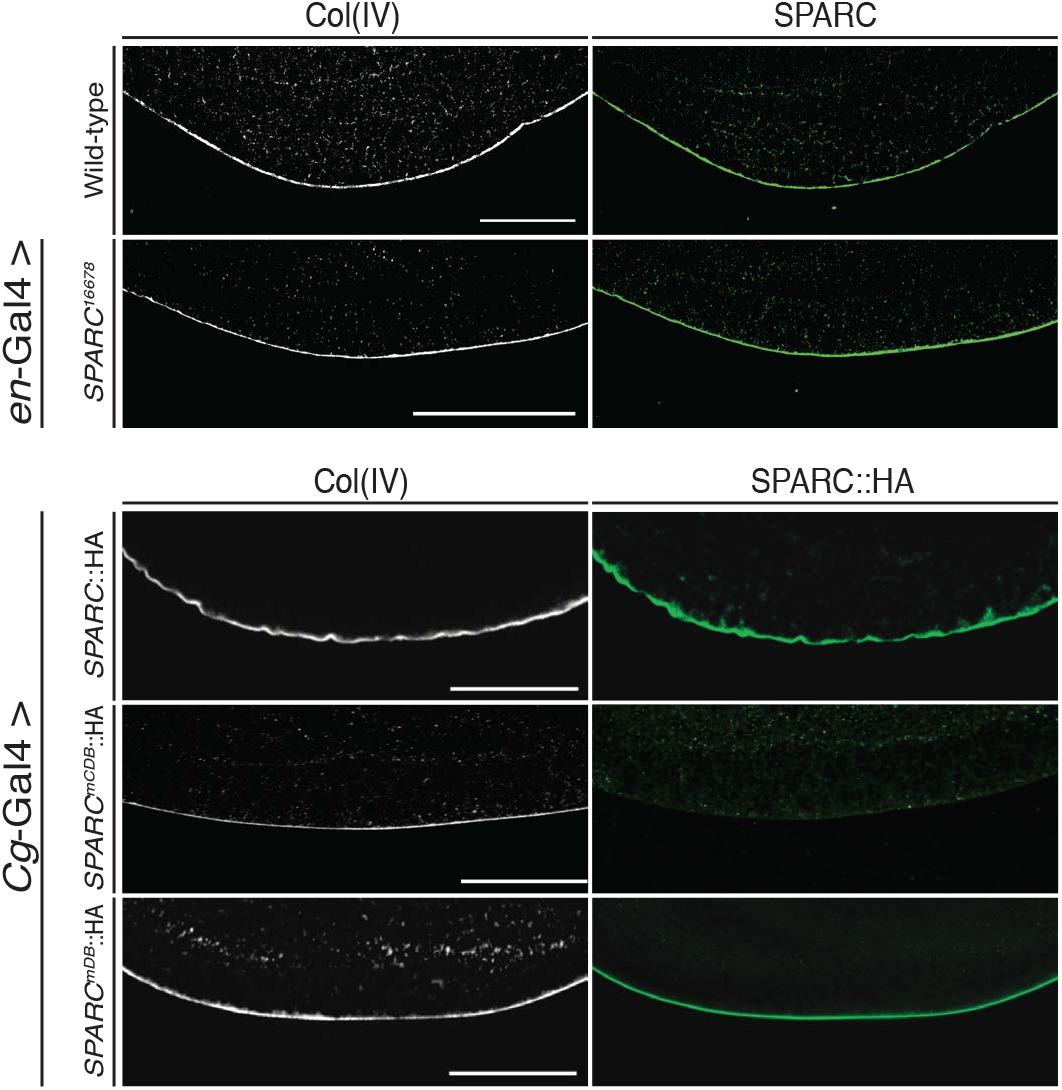
Disruption of the collagen-binding epitopes of SPARC, but not the disulfide bridge of EF-hand2 prevented localization of SPARC to the wing imaginal disc. HA Immunostaining of *Cg*-Gal4>*SPARC∷*HA and Cg-Gal4>*SPARC*^*mDB*^∷HA in a wild-type background showed fat body-derived *SPARC*∷HA localization in the BM of the wing imaginal disc, mirroring the immunostaining with anti-SPARC antibody. Likewise, knockdown of SPARC expression with *en*-Gal4 (*en*-Gal4, UAS-RFP>*SPARC*^*16678*^) did not impact the presence of SPARC or Col(IV) in the wing imaginal disc BM. In contrast, HA immunostaining of *Cg*-Gal4>*SPARC*^*mCBD*^∷HA in a wild-type background did not localize to the BM of the wing imaginal discs.

Studies in mammals have demonstrated tissue-specific differences in the affinity of SPARC for Col(IV) (Kelm and Mann, 1991). Since SPARC is produced by the wing imaginal disc in addition to the fat body (Flybase, http://flybase.org/reports/FBgn0026562), we examined whether SPARC derived from these two tissues differed in their colocalization with Col(IV) in BMs. Knockdown of SPARC expression by the wing imaginal disc using *SPARC*^*16678*^ RNAi driven by engrailed-Gal4 (*en*-Gal4) did not impact the localization of SPARC and Col(IV) in the wing imaginal disc (Fig. 5), with flies surviving to adulthood with no overt phenotype. Thus, the data support that SPARC found in the wing imaginal disc BM is predominantly derived from fat body adipocytes.

To confirm that wing imaginal disc-derived SPARC does not localize to the wing imaginal disc BM, we drove expression of *SPARC*∷HA in the wing imaginal disc. Despite the presence of Col(IV) in the wing imaginal disc, no HA immunoreactive SPARC colocalized to the wing imaginal disc BM. Similarly, the expression of *SPARC*^*mDB*^∷HA and *SPARC*^*mCBD*^∷HA by the wing imaginal disc did not associate to the wing imaginal disc BM (Fig. 6A). Rather, *SPARC*∷HA expressed by the wing imaginal disc was found to associate pericellularly within the fat body, distinct from the localization of Col(IV). This was also seen with *SPARC*^*mCBD*^∷HA expressed by the wing imaginal disc (Fig. 6B-C). Fat body- and wing imaginal disc-derived SPARC migrated comparably in a polyacrylamide gel, indicating similar molecular weight (Fig. 6D). Given that fat body adipocytes are polyploid, whereas wing imaginal disc cells are diploid, the levels of SPARC generated by the *en*-Gal4 driver were markedly less than that by the *Cg*-Gal4 driver. The collective data indicate that, as in mammals, tissue-specific variants of SPARC that likely differ in their affinity for Col(IV) are expressed by *Drosophila*.

**Fig. 6.**
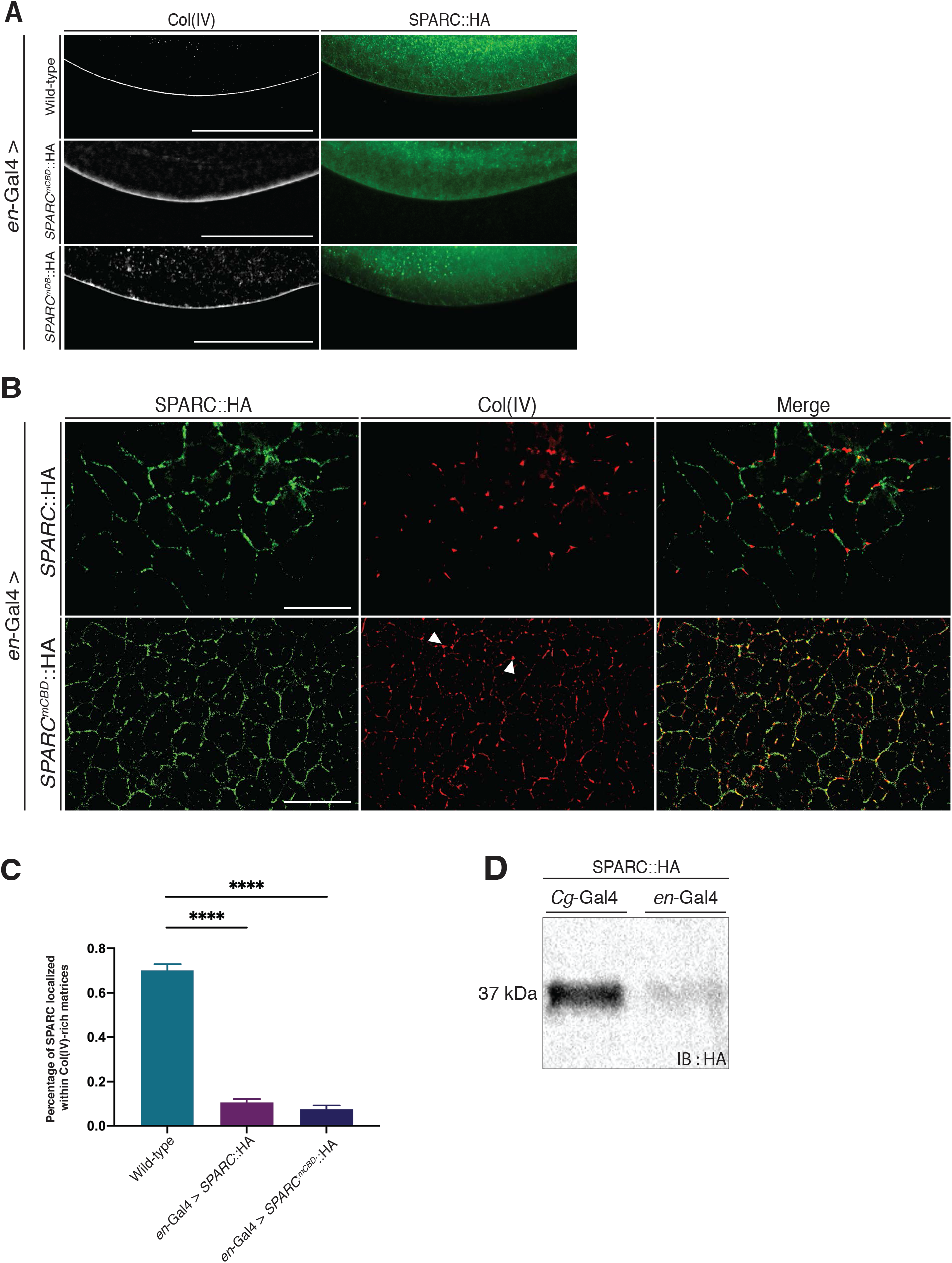
Wing imaginal disc-derived *SPARC*∷HA localizes to the fat body but does not colocalize with Col(IV). (**A**) HA immunostaining of wing imaginal disc-derived *SPARC*∷HA, *SPARC*^*mCBD*^∷HA and *SPARC*^*mDB*^∷HA in a wild-type background showed an absence of SPARC localization within the wing imaginal disc BM. Additionally, Col(IV) is present in the wing imaginal disc BM of all three genotypes. **(B)** HA immunostaining of 3^rd^ instar wing imaginal disc-derived *SPARC*∷HA, *SPARC*^*mCBD*^∷HA fat body in a wild type background revealed pericellular fat body adipocyte localization distinct from that of Col(IV). Arrowheads indicate CIVICs highlighted by immunostaining with anti-Col(IV) antibody. **(C)** Percentage of SPARC staining that colocalized with Col(IV) in wild-type, *en*-Gal4>*SPARC*^*mCBD*^∷HA and *en*-Gal4>*SPARC*^*mDB*^∷HA. For the latter two genotypes, anti-HA antibody was used. Data were analyzed by one-way ANOVA followed by Dunnett’s multiple comparisons test, P<0.0001 (****). **(E)** Western blot analysis showing levels of SPARC∷HA generated in 3^rd^ instar larvae by the *Cg*-Gal4 *vs. en*-Gal4 driver.

## Discussion

Previous studies have demonstrated a Col(IV) chaperone-like activity of SPARC in invertebrate model organisms (Isabella and Horne-Badovinac, 2015b; Morrissey et al., 2016; Shahab et al., 2015). In this study, we used a hemocyte cell line to determine that SPARC and Col(IV) first colocalize in the trans-Golgi with little or no colocalization within the ER or cis-Golgi. We confirmed, using a CRISPR-Cas9 approach, that disruption of the SPARC locus results in larval lethality at the 2^nd^ instar. We found that mutating two key amino acids in the Col(IV)-binding epitopes resulted in a similar larval lethality, demonstrating that the interaction of SPARC and Col(IV) is essential for survival. While the disulfide-bridged EF-hand2 of SPARC is known to enhance the binding of Col(IV) by the collagen-binding epitopes, this enhancement is not required for survival. Our data further indicate that mutation of the collagen-binding epitopes results in intracellular retention of Col(IV) within fat body adipocytes and a lack of Col(IV) within distal tissue BMs. In contrast, disruption of the disulfide-bridged EF-hand2 resulted in less pronounced intracellular retention with minimal impact on the diffusion of Col(IV) to distal tissue BMs. These findings indicate that the larval lethality due to the loss of SPARC expression results from the absence of Col(IV) chaperone-like activity.

An HA-tag was selected to track SPARC produced by specific tissues rather than a direct fluorescence tag. Preliminary studies tagging SPARC at its C-terminus with mCherry (UAS-*SPARC*∷mCherry) resulted in a non-functional fusion protein that was degraded *in vivo*. Furthermore, while not degraded, UAS-*SPARC*∷GFP was retained intracellularly and was therefore unable to rescue the loss of SPARC (Fig. S3B).

The ability of SPARC to colocalize with Col(IV) in BMs is dependent on its tissue of origin. Fat body-derived SPARC concentrates within BMs at both proximal and distal sites, whereas SPARC produced by the wing imaginal disc fails to associate with the BM of either the wing imaginal disc or fat body, despite the fact that it is able to diffuse to the latter. Moreover, the same cDNA was used to express SPARC in the wing imaginal disc as was used to express SPARC in the fat body. Thus, the difference in the behaviour of the SPARC protein generated by these two tissue sites can be attributed to post-translational processing and does not reflect a protein isoform.

A likely possibility for this finding is a difference in N-glycosylation, which has been shown to impact the ability of mammalian SPARC to bind to a variety of collagens (Kaufmann et al., 2004; Kelm and Mann, 1991). For example, bone-derived SPARC binds collagens whereas platelet-derived SPARC has no measurable binding to collagens, which has been attributed to their differences in N-glycosylation. We therefore postulate that wing imaginal disc-derived SPARC generates a SPARC glycoform that is distinct from fat body-derived SPARC and is unable to bind to Col(IV). It is important to note that the use of *en*-Gal4 to drive SPARC expression from the wing imaginal disc results in markedly less protein expression than SPARC driven in the fat body by *Cg*-Gal4 due to the presence of diploid cells in the wing imaginal disc *vs.* polyploid cells in the fat body. However, despite this difference in expression levels, SPARC generated from the wing imaginal disc was not detected locally and only detected in the fat body, distinct from Col(IV) localization. The function of wing imaginal disc-derived SPARC remains to be determined.

Fat body adipocytes are the principal source of BM components during *Drosophila* larval development. SEM micrographs indicate that funnel-shaped pores of variable diameter are randomly distributed over the surface of adipocyte BMs as we previously reported (Shahab et al., 2015). BM pores have been shown in several vertebrate tissues and have been hypothesized to have diverse functions, such as serving as a conduit for immune cell and growth factor trafficking, and the transmigration of extracellular vesicles (Howat et al., 2001; McAlindon et al., 1998). Altering the expression of SPARC by RNAi knockdown or by mutating either the evolutionary conserved collagen binding epitopes or the disulfide-bridged EF-hand2 of SPARC led to a fibrotic-like accumulation of Col(IV) surrounding adipocytes and to a striking dysregulation in the number and size of the BM pores. An increase in pore size was observed for *SPARC*^*1510D*^, *SPARC*^*mCBD*^∷HA and *SPARC*^*mDB*^∷HA. In contrast to the smooth lateral edges of pores observed in 2^nd^ and 3^rd^ instar wild-type fat body, the pores associated with the SPARC-mutant genotypes have elevated circumferential edges and a fibrotic-like inner core. These changes observed in the surface topography of BM pores may influence the regulatory role of the fat body in bidirectional trafficking of cells and bioactive molecules and vesicles.

A regulatory role of SPARC for Col(IV) homeostasis was hypothesized to begin with an intracellular association of SPARC with Col(IV) prior to their secretion. A chaperone-like activity has been proposed, suggesting that upon being co-secreted, SPARC inhibits Col(IV) from binding to integrins, thereby adequately delaying the nucleation phase of Col(IV) polymerization to allow a subset of Col(IV) to migrate and assemble in the BM of distal tissues in 3^rd^ instar larvae (Chioran et al., 2017). We now report that compared to 3^rd^ instar wild-type larvae, where intracellular Col(IV)-positive punctae are not readily visible by immunohistochemistry, 2^nd^ instar adipocytes contain many intracellular Col(IV)-positive punctae. These Col(IV) punctae increased dramatically in size in the absence of SPARC. Small intracellular laminin punctae also became visible in the absence of SPARC; however, these did not colocalize with SPARC and were much smaller in diameter than Col(IV) and SPARC immunoreactive punctae. The intracellular accumulation of Col(IV) could be attributed to a lack of secretion or an increase in endocytosis. The latter does not appear to be the cause as both Col(IV) and *SPARC*^*mCBD*^∷HA are observed within these punctae. Since *SPARC*^*mCBD*^∷HA does not bind to Col(IV) and does not colocalize with BMs, simple entrapment during endocytosis is an unlikely cause of their colocalization in these punctae. An issue that remains to be addressed is the mechanism by which *SPARC*^*mCBD*^∷HA is able to colocalize with Col(IV) in intracellular punctae. It also remains to be resolved why Col(IV) is not concentrated in intracellular punctae in 3^rd^ instar fat body adipocytes with the disruption of SPARC. Immunostaining of 2^nd^ instar larval fat body suggests a greater level of Col(IV) and SPARC synthesis than in 3^rd^ instar larvae. It is thus possible that the large intracellular Col(IV) and SPARC punctae observed in 2^nd^ instar, but not in 3^rd^ instar, are due to stage-specific differences in their level of synthesis.

Biochemical studies with mammalian SPARC have reported that the disulfide bridge in EF-hand2 augments the binding affinity of SPARC for Ca^2+^, which in turn enhances collagen binding (Pottgiesser et al., 1994). Studies performed *in vitro* indicate that disruption of the disulfide bridge in EF-hand2 results in decreased SPARC protein secretion secondary to decreased SPARC protein production or stability (Pottgiesser et al., 1994). In contrast to these findings, our *in vivo* data indicate that protein production and secretion is not diminished by the loss of the disulfide bridge. Interestingly, transgenic expression of *SPARC*^*mDB*^∷HA rescued SPARC protein-null flies to adulthood. While the fat bodies of 3^rd^ instar *SPARC*^*mDB*^∷HA larvae displayed fibrotic-like accumulation of BM components similar to those observed in the fat bodies of 2^nd^ instar *SPARC*^*mCBD*^∷HA or *SPARC*^*1510D*^, the phenotype was less severe. As *SPARC*^*mCBD*^∷HA exhibits larval lethality, it can therefore be concluded that the severity of disruption to BM homeostasis is the underlying mechanism responsible for larval lethality.

Overexpression of SPARC results in the loss of 3^rd^ instar fat body CIVICs, resulting in decreased adipocyte cell-cell adhesion (Dai et al., 2017). Our data indicate that CIVICs are first detected as larvae undergo their transition from 2^nd^ to 3^rd^ instar, coincident with the gradual 67-fold increase in adipocyte size from early 2^nd^ to wandering 3^rd^ instar. Our ultrastructural data indicate that overexpressing SPARC led to minor changes in adipocyte morphology at 2^nd^ instar in contrast to disrupted adipocyte cell-cell adhesion at 3^rd^ instar. Thus, until late 2^nd^ instar, the presence of a BM appears sufficient to maintain adipocyte cell-cell adhesion. By the time adipocytes increase in size to the level observed in early 3^rd^ instar larvae, stability of cell-cell adhesion appears dependent on the presence of CIVICs.

Previous studies have provided novel insight into the role of SPARC in the regulation of Col(IV) supramolecular assembly, biomechanics and dynamics in BMs during development (Dai et al., 2017; Isabella and Horne-Badovinac, 2015b; Pastor-Pareja and Xu, 2011; Shahab et al., 2015). Our structure-function analyses presented here indicate a critical requirement for the collagen-binding epitopes and the disulfide bridge in EF-hand2 of SPARC for the proper assembly of Col(IV) in BMs of proximal and distal tissues. Fig. 7 presents a model summarizing the key findings from this study. SPARC first associates with Col(IV) within the cis-Golgi and colocalizes within secretory vesicles. Co-secretion of SPARC with Col(IV) delays nucleation and polymerization of Col(IV), enabling a subset of Col(IV) to translocate to distal sites, such as the wing imaginal disc, where it undergoes polymerization and incorporation into BMs. SPARC produced by distal sites, such as the wing imaginal disc, migrates to the fat body where it exerts unknown functions independent of Col(IV) association. In the absence of SPARC, Col(IV) accumulates intracellularly within secretory vesicles and in the fat body BM. The unavailability of fat body-derived SPARC prevents Col(IV) incorporation in BMs at distal sites, resulting in larval lethality. Putative differences in N-glycosylation may account for the lack of Col(IV) colocalization with wing imaginal disc-derived SPARC. A better understanding of the diverse contributions of these two distinct SPARC populations to BM supramolecular assembly and fat body function has important implications for the understanding of SPARC functions in vertebrates.

**Fig. 7.**
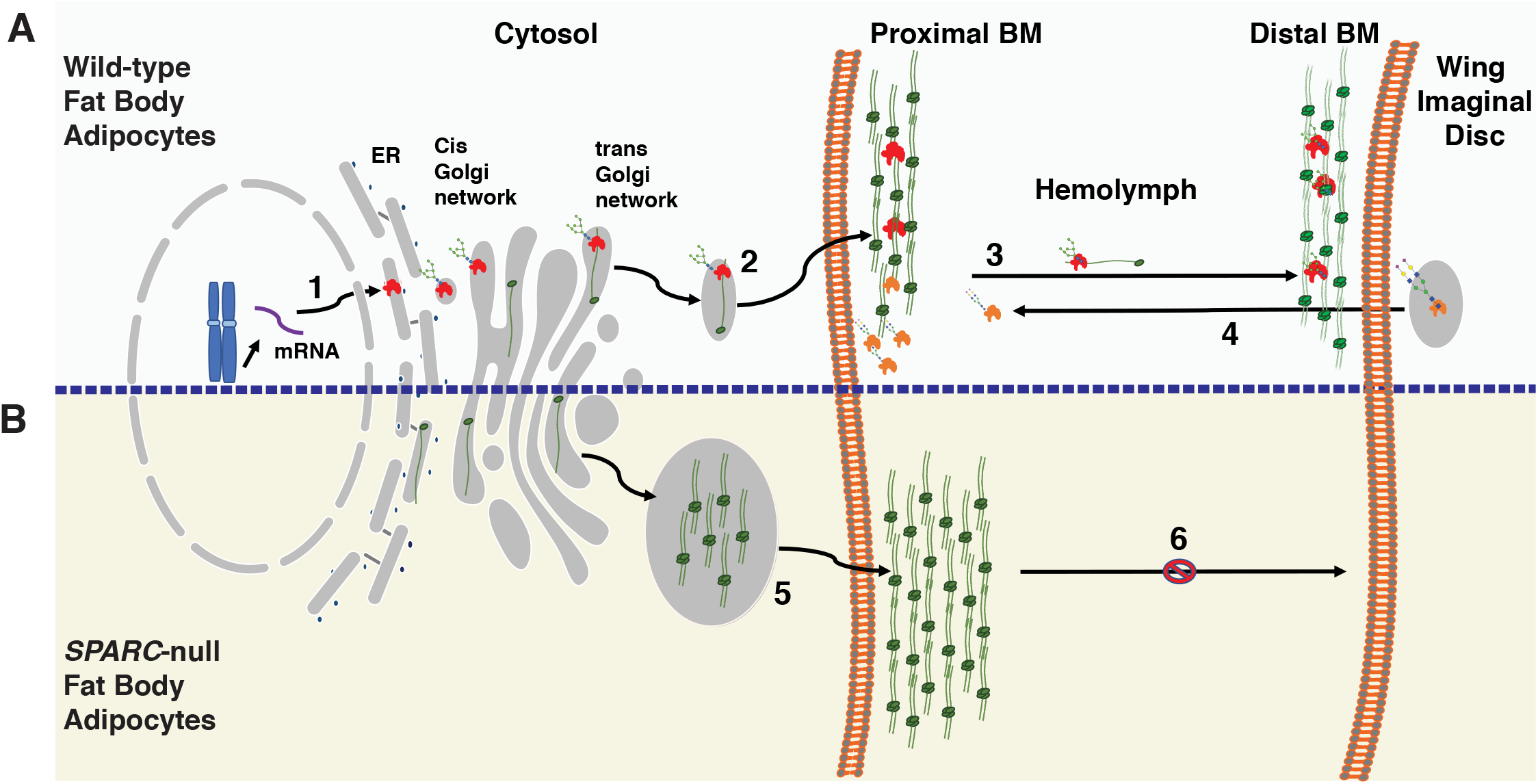
Model of two potential isoforms of SPARC, their localizations and interactions with Col(IV). **(A)** Sequence of events within wild-type larvae. **1.** Translation of SPARC within the ER and translocation to the trans-Golgi. **2.** SPARC and Col(IV) colocalize in the trans-Golgi just prior to their secretion. Fat body-derived SPARC (red) colocalizes with Col(IV) (green) in the fat body BM (proximal) and **3.** diffuses to and becomes incorporated within the wing imaginal disc BM (distal). **4.** In contrast, wing imaginal disc-derived SPARC (orange) diffuses to fat body cell borders where it does not colocalize with Col(IV). **(B)** Sequence of events SPARC protein-null larvae. **5&6.** In the absence of SPARC, Col(IV) is secreted by 2^nd^ and 3^rd^ instar fat body adipocytes, but forms a fibrotic-like network on the surface of the fat body adipocytes. The decrease in solubility of Col(IV) in the absence of SPARC may account for its inability to diffuse to distal BMs.

## Materials and Methods

### Generation of SPARC protein-null and mutant SPARC transgenic fly lines

Details on generation of the CRISPR-Cas9 SPARC protein-null fly lines are described in Fig. S1. cDNA for mutant SPARC constructs were synthesized by ThermoFisher Scientific using GeneArt Strings™ technology and inserted into pENTR 2B Dual selection vector from ThermoFisher Scientific at a 3:1 (insert:destination) vector molar ratio. The resultant entry vectors containing the desired SPARC constructs were recombined with the pPWH destination vector containing a UAS promoter and a C-terminus 3xHA tag. Expression vectors were injected into w^1118^, yw embryos for integration into the 2^nd^ chromosome attp16 locus by BestGene Inc. (Chino Hills, California). Flies containing the construct of interest were selected for the y^+^ marker (orange eyes) and were crossed with a double-balancer line (Bl/Cyo;TM2/TM6B) to establish the line.

### Western blot analyses

Tissues were homogenized with a pestle in microcentrifuge tubes in lysis buffer (100 mM tris 4% SDS pH 6.8). Protein concentration was measured using a Pierce BCA Protein Assay Kit (Pierce, Rockford, IL). 5-20 μg of each sample were diluted in 2X Laemmli buffer (100mM Tris pH 6.8, 200mM DTT, 20% glycerol, 0.01% Bromophenol blue and 4% SDS), boiled for 5 min, loaded into a 4-20% SDS polyacrylamide gel. Electrophoresed proteins were transferred to PVDF membranes. Membranes were blocked in 5% skim milk in PBST (PBS, containing 0.1% Tween 20) for 1 hr at room temperate prior to incubation with their appropriate primary antibodies listed in Table S1 at 4°C overnight. Membranes were washed 3 times with PBST for 10 min. Membranes were then incubated with their appropriate peroxidase-conjugated secondary antibodies listed in Table S1 for 1 hr at room temperature. Membranes were exposed to chemiluminescent reagents (Santa Cruz Catalog #SC-2048) and visualized using BioRad Molecular Imager Gel Doc™ XR+ Imaging System.

### Isolation of 2^nd^ and 3^rd^ instar fat body

Second and third instar larvae were placed in a dissecting well containing PBS. Forceps were used to pinch open the cuticle at the posterior end of the larvae. The anterior end was then held in place with forceps and the contents of the larvae were expelled from the cuticle by sliding the forceps in an anterior to posterior direction. Fat body from 3^rd^ instar larvae were then detached from other tissues. Due to the small size of 2^nd^ instar larvae, the fat body was not detached from other tissues prior to preparation for immunohistochemistry and SEM.

### Immunohistochemistry

Tissue samples were fixed in 4% formaldehyde in PBS for 45 min. Samples were then washed 3 times 20 min in PBSTx (PBS, 0.1% Triton X-100), blocked with 5% Donkey serum for 30 min, and then incubated with primary antibody in Donkey serum at dilutions indicated in Table S1 overnight at 4°C. Tissues were washed and blocked again in Donkey serum for 1 hr. Host-specific Alexa Fluor-conjugated secondary antibodies were added to the solution and incubated for an additional 2 hr. Tissue samples were washed 3 times in PBSTx before mounting on a slide coated with Vector shield mounting medium (Burlingame, California). All preparations were imaged by confocal microscopy (Leica TCS SP8 non-resonant) or spinning disc microscopy (Nikon Ti2 equipped with Yokogawa CSU-X1 head and Photometrics Myo CCD camera). Image analyses was performed using ImageJ software.

### SPARC lethality assessment

To assess larval lethality, rescue analyses were performed by crossing specific SPARC mutant flies with SPARC protein-null mutant flies (*SPARC*^*1510D*^). Crosses were carried out in cages and fly embryos were laid on grape juice-supplemented agar plates. The stage at which larval lethality occurred was scored based on the size of the larvae mouth hooks and shape of their spiracles.

### Measuring fat body adipocyte growth during larval development

The size of adipocytes was calculated using ImageJ by manually outlining the cell borders of adipocytes from early 2^nd^ and wandering 3^rd^ instar fat body images. Measurements were used to calculate the change in fat body adipocyte size. The average size of 2^nd^ and 3^rd^ instar fat body adipocytes were then compared to one another for statistical significance analysis using an unpaired t-test in GraphPad Prism^®^.

### Scanning electron microscopy (SEM)

Fat bodies from 2^nd^ and 3^rd^ instar larvae were dissected in PBS as described and were washed in 3% Glutaraldehyde, 0.1 M Sorensen’s Phosphate Buffer at 4°C for 24 hrs. The fixative was removed by washing tissues 3 times 10 min in 0.1 M Sorensen’s Phosphate Buffer. Tissues were then fixed in 1% osmium tetroxide, 0.1 M Sorensen’s Phosphate Buffer for 1.5 hrs and then washed 3 times for 10 min in 0.1 M Sorensen’s Phosphate Buffer. The tissues were dehydrated through an ascending ethanol series by incubating in 30-100% ethanol solutions for 10 min each and later infiltrated with an ethanol and hexamethyldisilizane (HMDS) series at ratio of 3:1, 1:1, 1:3 and 100% HMDS, all for 15 min each. Tissues were left to dry overnight in a depressed dish in 100% HMDS and later mounted on a specimen step. Samples were sputter coated with platinum using the Bal-Tec SCD050 and examined with a Hitachi S◻2500 scanning electron microscope at 1700X and 3700X magnification.

### Statistical analyses for the pore sizes and number of pores

Analysis of the pore sizes and number of pores were conducted by measuring their diameter using the respective scale bar and by counting the number of pores per sample via a 10.0 μm scale. For each sample, a total of five SEM images were used. Each sample comprised a total area of 1 mm2 at 3700X magnification. Statistical significance of the data was assessed by, one-way ANOVA followed by Dunnett’s multiple comparisons test (GraphPad Prism^®^, version 8.1.1).

### Fly stocks

All stocks and genotypes used in this study can be found in Table S2. All crosses were carried out at 25°C in standard vials.

## Acknowledgements

This work was supported by Discovery Grant 498474 from the Natural Sciences and Engineering Council of Canada to Dr. Maurice Ringuette. Cell lines were obtained from the Drosophila Genomics Resource Center, which is supported by NIH grant 2P40OD010949.

## Supplementary Figures

**Fig. S1. Generation and characterization of two CRISPR-generated SPARC mutants.**

**(A)** Schematic of the *Drosophila* SPARC gene locus (3R:26,869,238…26,871,995) and upstream and downstream adjacent genes. Exons are represented as boxes and the coding sequence is shaded in color. Deficiency lines used for complementation assays: the *Df(3R)BSC524* allele lacks genomic DNA between cytological bands 97C3 and 97D11; the *Df(3R)nm136* deletion (1963 bp) previously generated by P-element excision, is located within the His2Av gene adjacent to *SPARC*, but inhibits *SPARC* expression (Shahab et al., 2015). **(B)** A genomic map of the 3^rd^ chromosome encompassing the *SPARC* locus and the target sites and sequences used for sgRNA plasmid generation are indicated, with PAM sequences bolded and italicized. **(C)** DNA sequencing of PCR amplification of two CRISPR mutant line (*SPARC*^*1510B*^ and *SPARC*^*1510D*^) revealed 2761 bp deletions, represented by the red bar, spanning the *SPARC* locus and the genomic region encoding the 3’UTR of *Rb97D* (3R:26,870,690…26,873,451). **(D)** Complementation tests performed with the *Df(3R)BSC524* deficiency line and potential CRISPR-generated mutants. Each solid line represents the percentage of lethal progeny in each stage of development from the mating of a single male from subgroups 1 through 4 with a female carrying the *Df(3R)BSC524* allele. Each dotted line represents the percentage of survivors from the same collection of progeny. Progeny from four males in CRISPR subgroup 1 (1A, 1B, 1C, 1D) could not be complemented by the *Df(3R)BSC524* allele and exhibited larval lethality primarily at 2^nd^ instar, indicative of a genetic alteration on the right arm of the third chromosome. **(D)** Larval lethality assay results for the two CRISPR-generated allele, referred to as *SPARC*^*1510B*^ and *SPARC*^*1510D*^. Whether in trans to *Df(3R)nm136, H2Av*∷*GFP* or homozygous for the two CRISPR-generated alleles, all show larval lethality primarily at 2^nd^ instar.

**Fig S2. Molecular characterization of the *SPARC***^***1510D***^ **mutant**.

**(A)** *SPARC*^*1510D*^ homozygous embryos were generated by crossing *SPARC^1510D^/ TM6B, Sb, Dfd-YFP* males and females. The absence of the YFP in the subesophageal ganglion was used to differentiate *SPARC*^*1510D*^ homozygous embryos from *SPARC*^*1510D*^/ *TM6B, Sb, Dfd-YFP* embryos within the total collection of progeny. Stage 15-16 *SPARC*^*1510D*^ homozygous embryos were immunostained with anti-SPARC and anti-Col(IV) antibodies and counterstained with DAPI to visualize chromatin. The arrow in the enlarged image of the merged image indicate Col(IV) expression in circulating hemocyte cells whereas the arrowhead indicate normal deposition of Col(IV) into BMs. **(B)** Stage 15-16 *SPARC*^*1510B*^/*TM6B, Sb, Dfd-YFP* embryos were immunostained with anti-SPARC and anti-Col(IV) antibodies were used as a positive control for SPARC expression. Yellow Fluorescence Protein (YFP) detection in the subesophageal ganglion confirms the presence of the TM6B balancer allele. Enlarged images of the boxed region highlight the colocalization of SPARC and Col(IV) in circulating hemocytes (arrow) and colocalization of SPARC and Col(IV) in the BM (arrowhead).

**Fig S3. *SPARC*∷GFP protein is not degraded, but is retained intracellularly**.

**(A)** Western blot analysis of *SPARC*∷HA lysate from 3^rd^ instar larvae immunoblotted with anti-SPARC and anti-GFP antibodies indicating immunoreactive bands at the expected molecular weights. **(B)** Wild-type 3^rd^ instar fat body immunostaining with anti-SPARC antibody reveals pericellular (white arrow) localization while immunostaining with anti-GFP reveals intracellular retention of *SPARC*∷GFP (yellow arrow) and no staining observed extracellularly.

**Fig S4. SEM analysis overview**.

**(A)** Table depicting the genotype, larval growth stage (2nd or 3rd instar), average pore size (in μm), average number of pores (per mm^2^), and standard error of the mean. Graphical presentation of the mean number of pores for each genotype at 2^nd^ **(B)** and 3^rd^ **(C)** instar. Graphical presentation of the mean pore diameter for each genotype at 2^nd^ **(D)** and 3^rd^ **(E)** instar in relation to the mean size of pores.

**Table S1. List of antibodies used and their dilution and source.**

**Table S2. List of fly lines used in this study, their description and source.**

